# Trade-offs between host tolerances to different pathogens in plant-virus interactions

**DOI:** 10.1101/799007

**Authors:** Nuria Montes, Viji Vijayan, Israel Pagán

## Abstract

Although accumulating evidence indicates that tolerance is a plant defence strategy against pathogens as widespread as resistance, how plants evolve tolerance is poorly understood. Theory predicts that hosts will evolve to maximize tolerance or resistance, but not both. Remarkably, most experimental works failed in finding this trade-off. We tested the hypothesis that the evolution of tolerance to one virus is traded-off against tolerance to others, rather than against resistance, and identified the associated mechanisms. To do so, we challenged eighteen *Arabidopsis thaliana* genotypes with *Turnip mosaic virus* (TuMV) and *Cucumber mosaic virus* (CMV). We characterized plant life-history trait modifications associated with reduced effects of TuMV and CMV on plant seed production (fecundity tolerance) and life period (mortality tolerance), both measured as a norm of reaction across viral loads (range tolerance). Also, we analysed resistance-tolerance and tolerance-tolerance trade-offs. Results indicate that tolerance to TuMV is associated with changes in the length of the pre-reproductive and reproductive periods, and tolerance to CMV with resource reallocation from growth to reproduction; and that tolerance to TuMV is traded-off against tolerance to CMV in a virulence-dependent manner. Thus, this work provides novel insights on the mechanisms of plant tolerance and highlights the importance of considering the combined effect of different pathogens to understand how plant defences evolve.

## Introduction

Parasitism is a lifestyle chosen by 50% of all known organisms (*Poulin & Morand, 2000*). This means that, along their lifespan, hosts will be recurrently challenged by parasites. Parasites may be pathogens, causing diseases that have a negative impact on the fitness of infected hosts, i.e., virulence (*Read, 1994; Anderson et al., 2004*). To cope with pathogens, hosts have developed a variety of defence mechanisms to avoid/limit infection and its negative effects (*Agnew et al., 2000*), which have relevant consequences for the fitness of both interacting partners (*Woolhouse et al., 2002*). Thus, investigating the evolution and the mechanistic basis of these defences is central to understand the dynamics of host-pathogen interactions (*Jones & Dangl, 2006; Pagán & García-Arenal, 2018*).

The two main host defences against pathogens are resistance, i.e., the host’s ability to limit pathogen multiplication (*Clarke 1986; Strauss & Agrawal, 1999*), and tolerance, i.e., the host’s ability to reduce the effect of infection on its fitness at a given pathogen load (*Little et al. 2010; Råberg, 2014*). They represent two different strategies to deal with pathogens: resistance reduces the risk of infection and the multiplication rate of the pathogen, whereas tolerance does not. Hence, it is predicted that if hosts evolve resistance the prevalence of the pathogen in the host population will decrease, whereas tolerance will increase prevalence (*Roy & Kirchner, 2000*). Consequently, both resistance and tolerance may have significant, but different, impact on the dynamics of host and pathogen populations (*Roy & Kirchner, 2000; Pagán & García-Arenal, 2018*). Researchers have devoted considerable effort to understand the molecular basis and evolutionary consequences of resistance to pathogens. However, tolerance has received comparatively less attention, and the processes shaping its evolution are only partially understood (*Little et al., 2010; Pagán & García-Arenal, 2018*).

A body of mathematical work has modelled the conditions in which tolerance evolves. Early models assumed that resources are limited and can be diverted into resistance or tolerance, but not both, and predicted that tolerance or resistance would prevail because they were mutually exclusive (*van der Meijden et al., 1988; Herms & Mattson, 1992*). More recent models incorporated the idea that resistance and tolerance might not be fully exchangeable, and predicted that both defence mechanisms would coexist, with host fitness maximized: (i) only at maximum tolerance or maximum resistance (*Mauricio et al., 1997; Boots & Bowers, 1999*), or (ii) at intermediate levels of both (*Restif & Koella, 2003,2004; Fornoni et al., 2004*). In none of these scenarios tolerance and resistance can be maximized simultaneously. Hence, the common idea underlying the theory on the evolution of tolerance is that there is a trade-off between resistance and tolerance. However, there is remarkably little experimental support for such trade-off in host-pathogen, and particularly in plant-pathogen, interactions. Indeed, most studies on plant viruses (*Carr et al., 2006; Pagán et al., 2007,2009; Montes et al., 2019*), bacteria (*Kover & Schaal, 2002; Goss & Bergelson, 2006*) and fungi (*Simms & Triplett, 1994*) failed in finding a resistance-to-tolerance negative association.

A possible explanation for this lack of support of a resistance-tolerance trade-off is that other forces might come into play in shaping the evolution of plant defences. In nature, plant populations are challenged by multiple pathogens (*Syller, 2012*), not necessarily coinfecting the same individuals, and the evolution of tolerance to one pathogen may depend on the interaction with tolerances to others. According to the Life-History Theory, hosts may achieve tolerance to pathogens through modifications of their life-history (*Minchella, 1985; Stearns, 1992*). These changes may respond to two contrasting mechanisms: Highly virulent pathogens will induce shorter host pre-reproductive, and longer reproductive, periods in order to produce progeny before resource depletion, castration or death. Conversely, less virulent pathogens will induce host resource reallocation from growth to reproduction, and/or a delay in host reproduction, which would allow compensating the pathogen effect on host fitness (*Hochberg et al., 1992; Gandon et al., 2002*). Hence, depending on the pathogen’s virulence, tolerance may require markedly different, even opposed, host responses that likely are difficult to maximize simultaneously. As a consequence, trade-offs between tolerances to different pathogens might be important forces for the evolution of plant defences (Figure 1). Interestingly, such trade-offs have seldom been considered nor in mathematical models, or in experimental analysis (*Kutzer & Armitage, 2016; Pagán & García-Arenal 2018*).

**Figure 1.**
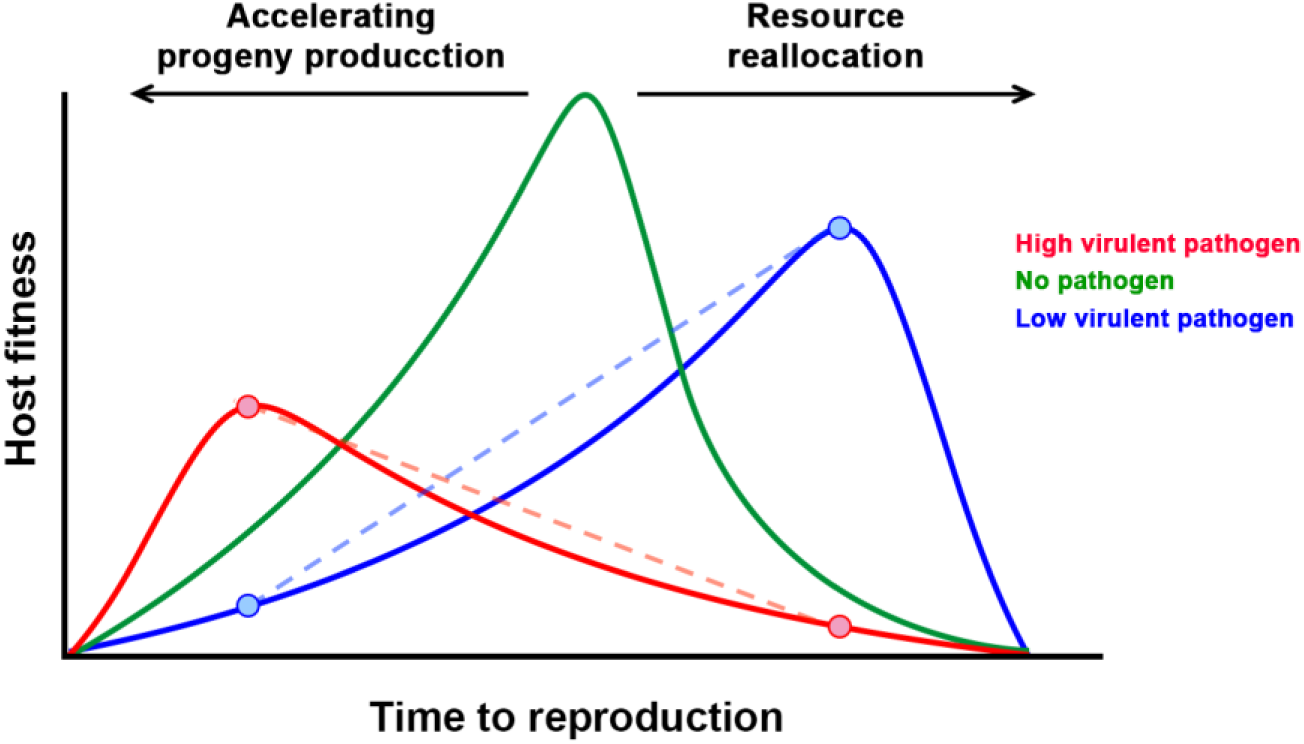
Which way to go? According to the Life-history Theory, hosts would modify their time to reproduction in opposite ways in order to achieve tolerance: when infected by a highly virulent pathogen (red line), hosts would bring forward reproduction to produce progeny before death; and when infected by a low virulent pathogen (blue line), host would delay reproduction so they can reallocate resourced from growth to reproduction. These strategies would maximize fitness in the presence of one virus at the cost of reducing fitness in the presence of the other (crossed dotted lines), establishing a tolerance-tolerance trade-off.

To address this central question to understand how plant defences against pathogens evolve, we utilized *Turnip mosaic virus* (TuMV, *Potyviridae*) and *Cucumber mosaic virus* (CMV, *Bromoviridae*), and *Arabidopsis thaliana* (from here on “Arabidopsis”, Brassicaceae). Both viruses are commonly found in wild populations of Arabidopsis at up to 80% prevalence (*Pagán et al., 2010*), indicating that the Arabidopsis–TuMV and Arabidopsis–CMV interactions are significant in nature. CMV infection moderately reduces seed production, rarely inducing sterility, and has little effect on plant life period (*Pagán et al., 2007,2008; Hily et al., 2016; Montes et al., 2019*). Thus, CMV can be considered as a moderately virulent virus. On the other hand, TuMV infection affects Arabidopsis flower and silique viability, which may severely affect plant fertility and often leads to sterility (*Sánchez et al., 2015*). Moreover, this virus greatly reduces plant life period (*Vijayan et al., 2017*). Therefore, TuMV can be regarded as a highly virulent pathogen in Arabidopsis, although milder TuMV genotypes exist (*Sánchez et al., 2015*). Interestingly, although both viruses have high prevalence and share common vectors (e.g., *Fujisawa, 1985*), in Arabidopsis CMV+TuMV mixed infections occurred at low frequency (*Pagán et al., 2010*), opening the possibility of evolving different tolerance responses to these two viruses that vary in virulence. Tolerance to CMV varies across Arabidopsis genotypes as a quantitative trait; and long-lived genotypes with low seed production to total biomass ratio (Group 1 genotypes) are generally more tolerant than short-lived genotypes that have high seed to biomass ratio (Group 2 genotypes) (*Pagán et al., 2008; Hily et al., 2016*). Tolerance to CMV in Group 1 genotypes is attained through modifications of life-history traits, mainly the reallocation of resources from growth to reproduction and, to a lesser extent, elongation of the pre-reproductive period (*Pagán et al., 2008; Shuckla et al., 2018*). Virus-induced resource reallocation appears to be CMV-specific, and it is not triggered upon TuMV infection (*Shuckla et al., 2018*). However, these authors used a reduced set of Arabidopsis genotypes, and did not test virulence-specific modifications of other life-history traits that would confer tolerance, and their potential trade-offs.

The key variables for measuring tolerance may vary depending on each plant-pathogen interaction (*Day, 2002; Rohr et al., 2010*). For instance, pathogens may affect plant fecundity directly or through reducing survival. In plants infected by a sterilizing pathogen such as TuMV, enhanced survival may represent the difference between reproducing or dying during the growth period. Thus, considering both the effect of infection on plant progeny production (fecundity tolerance) and survival (mortality tolerance) may be equally important to understand the evolution of tolerance. Conversely, plant mortality tolerance might be less relevant upon infection with a milder pathogen such as CMV, as infected plants generally reach the adult stage and reproduce. However, in most experimental analyses of tolerance to plant pathogens host fitness was measured only as progeny production (*Pagán & García-Arenal, 2018*), and the relationship between fecundity tolerance and mortality tolerance have been seldom analysed *(Pagán et al., 2008; Shuckla et al., 2018*). Another point under debate in the literature on plant tolerance is how it is quantified. Most often, tolerance has been measured as the effect of infection at a given pathogen load (i.e., point tolerance) (*Pagán & García-Arenal, 2018*). At odds, it has been proposed that a more informative approach is quantifying tolerance as the slope of a regression of host fitness against pathogen load (i.e., range tolerance); the steeper the slope, the lower the tolerance, which cannot be measured on a single plant but across individuals of a given host type (e.g. genotype) (*Little et al., 2010; Kutzer & Armitage, 2016*). Notably, range tolerance to plant pathogens has been seldom analysed to date (*Pagán & García-Arenal, 2018*).

Herein, we analyse whether Arabidopsis achieves (range) tolerance to TuMV infection and if such tolerance is related to modifications of plant life-history traits. Specifically, we analysed the association between the effect of infection on plant progeny production (fecundity tolerance) and life period (mortality tolerance) with resource reallocation from growth to reproduction and with modifications in the length of the growth and reproductive periods. We also analysed resistance-tolerance trade-offs upon infection by TuMV and CMV, and if tolerance to TuMV is traded-off against tolerance to CMV.

## Results

### Virus multiplication in Arabidopsis

The level of UK1-TuMV, LS-CMV and JPN1-TuMV RNA accumulation was used to evaluate Arabidopsis resistance to virus infection. Accumulation differed according to the virus (Wald *χ*^*2*^_2,448_ =211.52, *P*=1×10^−4^), and the interaction between virus and host genotype was significant (Wald *χ*^*2*^_34,448_=475.28, *P*<1×10^−4^). Thus, we analysed accumulation for each virus separately, considering plant genotype and allometric group as factors. For all three viruses, accumulation significantly differed between Arabidopsis genotypes (Wald *χ*^*2*^≥137.17, *P*<1×10^−4^), but not between allometric groups (Wald *χ*^*2*^≤0.41, *P*≥0.524) (Supplementary Table S1). Thus, the allometric group did not affect the level of resistance.

Broad-sense heritability of virus accumulation ranged from moderate to high depending on the virus: *h*^*2*^ _*b*_ =0.43, 0.60 and 0.68, for UK1-TuMV, LS-CMV and JPN1-TuMV, respectively (Supplementary Table S2). Therefore, there is significant genetic variation among the studied Arabidopsis genotypes for the ability to sustain virus multiplication.

### Arabidopsis fecundity and mortality tolerance to virus infection

Fecundity and mortality tolerances (slopes of the *SW* and *LP* to virus accumulation regression, respectively) differed depending on the virus (Wald *χ*^*2*^_2,48_≥143.28, *P*≤1×10^−4^). Both tolerance measures were smallest to UK1-TuMV and greatest to LS-CMV, with tolerances to JPN1-TuMV showing intermediate values (Figure 2 and Supplementary Table S1). The interaction between virus and Arabidopsis allometric group was also significant (Wald *χ*^*2*^_2,48_≥24.36, *P*≤1×10^−4^). Thus, fecundity and mortality tolerances were analysed for each virus independently. From here on, results will be presented firstly for the two viruses at the tolerance extremes (UK1-TuMV and LS-CMV), and lastly for the intermediate state (JPN1-TuMV).

**Figure 2.**
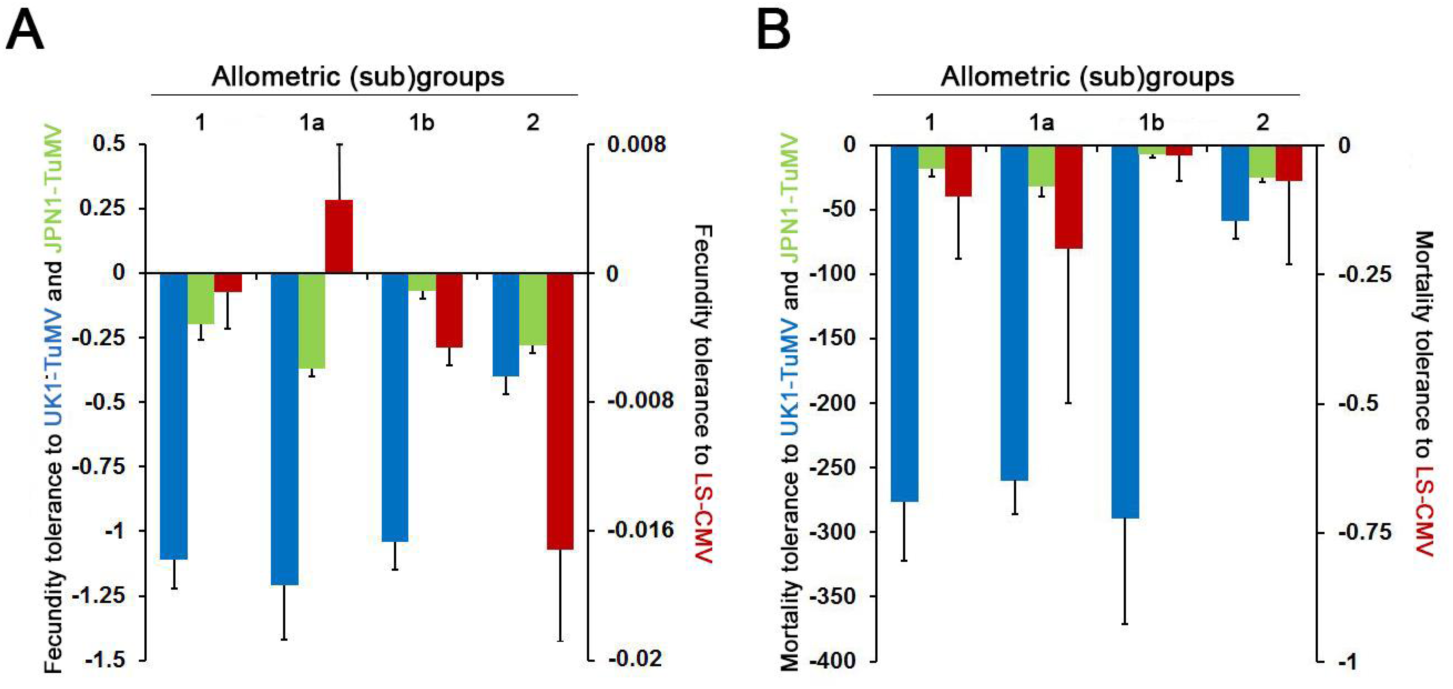
Arabidopsis fecundity and mortality tolerance to UK1-TuMV, LS-CMV and JPN1-TuMV. Panel A: Values of fecundity tolerance to UK1-TuMV (blue), to JPN1-TuMV (green) and to LS-CMV (red) measured as the slope of the *SW* to virus accumulation linear regression. Panel B: Values of mortality tolerance to the same three viruses measured as the slope of the *LP* to virus accumulation linear regression. Data are presented for allometric groups 1 and 2, and for subgroups 1a and 1b, and are mean ± standard errors across plant genotypes.

In general, viruses significantly reduced Arabidopsis fecundity (*SW*_*i*_*/SW*_*m*_<1) (Figure 3). Exceptions were LS-CMV-infected Cum-0 and Ll-0 plants (Group 1) that overcompensated the effect of virus infection (*SW*_*i*_*/SW*_*m*_>1) (Supplementary Table S1). Upon UK1-TuMV infection, fecundity tolerance varied according to the allometric group (Wald *χ*^*2*^_*1,16*_= 23.68, *P*<1×10^−4^). The negative slope was stepper for Group 1 genotypes, with none of the infected plants producing seeds, than for Group 2 ones, with only 61.8% of sterilized plants and fertile individuals in 8/11 genotypes (Figure 2 and Supplementary Table S1). Thus, Group 2 plants had higher fecundity tolerance. All LS-CMV-infected plants were fertile, with stepper negative slopes of the *SW* to virus accumulation regression for Group 2 genotypes (lower fecundity tolerance) than for Group 1 ones (Wald *χ*^*2*^_*1,16*_= 12.34, *P*<1×10^−4^). At odds with UK1-TuMV and LS-CMV infected plants, in JPN1-TuMV-infected plants the slope of the *SW* to virus accumulation regression did not differ between allometric groups (Wald *χ*^*2*^_*1,16*_=0.83, *P*=0.362). However, Group 1 genotypes showed a bimodal response: In Cum-0, Kas-0 and Ll-0 (Subgroup 1a), 50-70% of infected individuals were sterilized and regression slopes were steep. Conversely, infected Cad-0, Cdm-0, Kas-2 and Kyo-1 plants (Subgroup 1b) produced seeds and regression slope were shallower than for Subgroup 1a (Wald *χ*^*2*^_*1,5*_= 46.19, *P*<1×10^−4^) (Figure 2 and Supplementary Table S1). Group 2 genotypes, where 92% of infected plants produced seeds, showed intermediate and significantly different slope values than the two Group 1 subsets (Wald *χ*^*2*^≥ 4.61, *P*≤0.040) (Figure 2).

**Figure 3.**
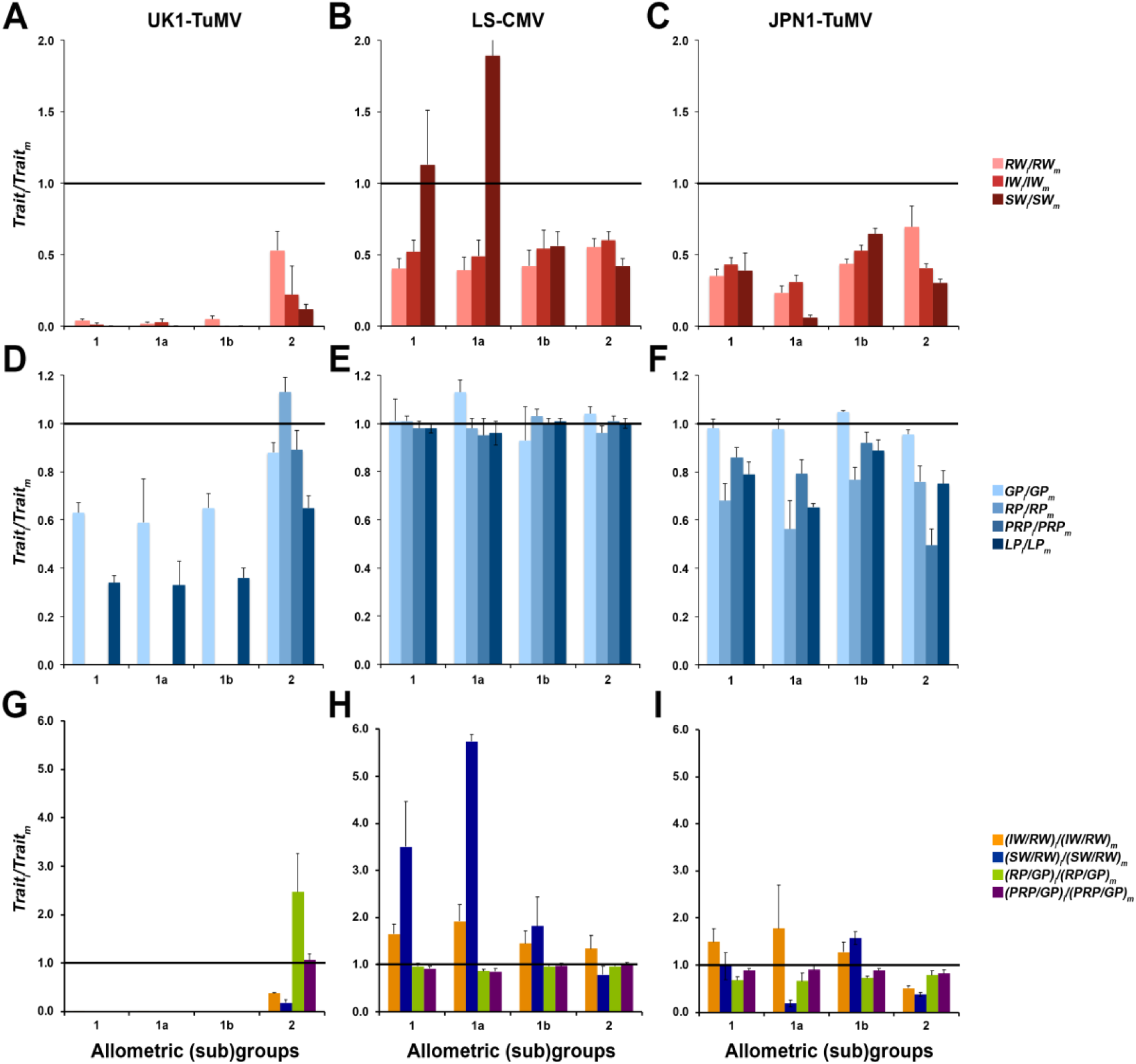
Effect of UK1-TuMV, LS-CMV and JPN1-TuMV infection on life-history traits for Arabidopsis allometric groups and subgroups. Panels A-C: Effect of viral infection on rosette weight (*RW*), inflorescence weight (*IW*) and seed weight (*SW*). Panels D-F: Effect of viral infection on growth period (*GP*), reproductive period (*RP*) and post-reproductive period (*PRP*). Panels G-I: Effect of infection on the ratios *IW/RW, SW/RW, RP/GP* and *PRP/GP*. All effects were estimated as the ratio between infected (i) and mock-inoculated (m) plants. Data are presented for allometric groups 1 and 2, and for subgroups 1a and 1b, and are mean ± standard errors of plant genotype means.

UK1-TuMV also reduced plant survival (*LP*_*i*_*/LP*_*m*_<1) (Figure 3), with mortality tolerance differing between allometric groups (Wald *χ*^*2*^_*1,16*_= 29.69, *P*=1×10^−4^). Mortality tolerance was smaller (steeper negative slope of the *LP* to virus accumulation regression) for Group 1 than for Group 2 plants (Wald *χ*^*2*^_*1,16*_= 29.69, *P*=1×10^−4^) (Figure 2). In contrast, LS-CMV infection had little effect on Arabidopsis survival: No differences between allometric groups were observed in the slope of the *LP* to virus accumulation regression (Wald *χ*^*2*^_*1,16*_= 0.02, *P*=0.900), indicating similar mortality tolerance, with very little effect of virus infection on *LP* as denoted by *LP*_*i*_*/LP*_*m*_ values near one (Figures 2 and 3). As for JPN1-TuMV-infected plants, mortality tolerance did not vary between allometric groups (Wald *χ*^*2*^_*1,16*_= 1.25, *P*=0.263). However, again Group 1 plants presented a bimodal response to infection: the slope of the *LP* to virus accumulation regression was significantly steeper in Subgroup 1a than in Subgroup 1b (Wald *χ*^*2*^_*1,5*_=14.25, *P*=1×10^−3^) and in Group 2 genotypes (Wald *χ*^*2*^_*1,13*_=8.34, *P*=0.004), which showed similar mortality tolerance (Wald *χ*^*2*^_*1,13*_=0.83, *P*=0.363), (Figure 2 and Supplementary Table S1). Note that upon infection by UK1-TuMV and LS-CMV Group 1 genotypes did not show this bimodal distribution (Wald *χ ;*^*2*^≤0.49, *P*≥0.483) (Supplementary Table S1).

Because fecundity and mortality tolerances are genotype-specific rather than plant-specific variables, by definition heritability for these traits could not be calculated.

### Relationship between modifications of Arabidopsis life-history traits and tolerance to virus infection

For each virus, the effect of infection on Arabidopsis growth and reproduction was quantified as the ratios of rosette, inflorescence and seed weights between infected and mock-inoculated plants (*RW*_*i*_*/RW*_*m*_, *IW*_*i*_*/IW*_*m*_ and *SW*_*i*_*/SW*_*m*_, respectively) (Figure 3A-C and Supplementary Table S1). In general, virus infection reduced *RW, IW* and *SW* (Wald *χ*^*2*^≥49.52; *P*<1×10^−4^), this reduction always depending on the Arabidopsis genotype (Wald *χ* ^*2*^≥388.79; *P*<1×10^−4^). For UK1-TuMV-infected plants, all ratios also depended on the allometric group (Wald *χ*^*2*^_*1,168*_≥25.87; *P*<1×10^−4^). In groups 1 and 2, *RW*_*i*_*/RW*_*m*_ was greater than *IW*_*i*_*/IW*_*m*_ (Wald *χ*^*2*^≥20.04; *P*<1×10^−4^) and *SW*_*i*_*/SW*_*m*_ (Wald *χ*^*2*^≥50.20; *P*<1×10^−4^) suggesting no resource reallocation from growth to reproduction. Indeed, (*IW/RW*)_i_/(*IW/RW*)_m_ and (*SW/RW*)_i_/(*SW/RW*)_m_ were always smaller than one (Wald *χ*^*2*^_*1,168*_≥21.35, *P*<1×10^−4^) (Figure 3G and Supplementary Table S1). The effect of LS-CMV on *RW* and *SW* (Wald *χ*^*2*^_*1,163*_≥4.33, *P*≤0.037), but not on *IW* (Wald *χ*^*2*^_*1,163*_=1.955, *P*=0.162), varied according to the allometric group. For Group 1, the effect of LS-CMV infection on *RW* was larger than on *SW* (Wald *χ*^*2*^_*1,49*_=10.89, *P*=1×10^−3^), whereas the opposite was observed in Group 2 (Wald *χ*^*2*^_*1,102*_=13.90, *P*=2×10^−4^). (*SW/RW)*_*i*_*/(SW/RW)*_*m*_ differed between allometric groups (Wald *χ*^*2*^_*1,163*_=11.77, *P*<1×10^−4^), values being greater than one only for Group 1 (Wald *χ*^*2*^_*1,60*_=7.11, *P*=0.008) (Figure 3H and Supplementary Table S1). Similar trends were observed in JPN1-TuMV-infected plants (Figure 3C), for which (*IW/RW)*_*i*_*/(IW/RW)*_*m*_ (Wald *χ*^*2*^_*1,161*_=2.66, *P*=0.003) and (*SW/RW)*_*i*_*/(SW/RW)*_*m*_ (Wald *χ*^*2*^_*1,161*_=17.18, *P*<1×10^−4^) were also greater for Group 1 than for Group 2 plants, values being similar or greater than one only for Group 1 (Figure 3I and Supplementary Table S1). These results would be compatible with resource reallocation from growth to reproduction in LS-CMV- and JPN1-TuMV-infected Group 1 plants. Again, JPN1-TuMV-infected Group 1 genotypes showed a bimodal distribution: *RW*_*i*_*/RW*_*m*_, *IW*_*i*_*/IW*_*m*_ and *SW*_*i*_*/SW*_*m*_ were smaller for Subgroup 1a than for Subgroup 1b (Wald *χ*^*2*^_*1,161*_≥5.65, *P*≤0.017) (Figure 3C), and the same was observed for (*SW/RW)*_*i*_*/(SW/RW)*_*m*_ (Wald *χ*^*2*^_*1,161*_=5.76, *P*=0.016) (Figure 3I). This ratio was greater than one only for Subgroup 1b genotypes (Wald *χ*^*2*^_*1,35*_=80.95, *P*<1×10^−4^), indicating that resource reallocation was associated with fecundity tolerance in this subgroup (Figure 3I and Supplementary Table S1).

We also quantified the effect of infection on the plant growth, reproductive and post-reproductive periods (*GP*_*i*_*/GP*_*m*_, *RP*_*i*_*/RP*_*m*_ and *PRP*_*i*_*/PRP*_*m*_, respectively) (Figure 3D-F and Supplementary Table S1). Upon UK1-TuMV infection, *GP*_*i*_*/GP*_*m*_ depended on the allometric group (Wald *χ*^*2*^_*1,166*_=17.95, *P*<1×10^−4^), this ratio being smaller for Group 1 than for Group 2 plants. Interestingly, in Group 2 genotypes the effect of infection on *GP* was greater than the effect on *RP* (Wald *χ*^*2*^_*1,39*_=52.46, *P*<1×10^−4^): *GP*_*i*_*/GP*_*m*_ was significantly smaller (Wald *χ*^*2*^_*1,39*_=9.73, *P*=0.002), and *RP*_*i*_*/RP*_*m*_ greater (Wald *χ*^*2*^_*1,39*_=7.55, *P*=0.006), than one. Thus, upon UK1-TuMV infection more tolerant Group 2 genotypes shortened their growth period but elongated the time dedicated to reproduction, as indicated by (*RP/GP)*_*i*_*/(RP/GP)*_*m*_ values greater than one in this subgroup (Figure 3G and Supplementary Table S1). Note that in Group 1 genotypes *RP* and *PRP* could not be quantified because plants did not produce mature siliques (Figure 3D and Material and Methods). On the other hand, LS-CMV infection did not affect *GP, RP* and *PRP* (Wald *χ*^*2*^_*1,166*_≤1.94, *P*≥0.164) their ratios being always near one in both allometric groups (Wald *χ*^*2*^≤0.76, *P*≥0.383) (Figure 3E). Accordingly, (*RP/GP)*_*i*_*/(RP/GP)*_*m*_ and (*PRP/GP)*_*i*_*/(PRP/GP)*_*m*_ were also near one (Wald *χ*^*2*^≤2.47, *P*≥0.116) (Figure 3H and Supplementary Table S1). Exception to this rule was Subgroup 1a, which included Arabidopsis genotypes that overcompensated the effect of LS-CMV infection on *SW*. These genotypes significantly elongated *GP*, as indicated by *GP*_*i*_*/GP*_*m*_ values higher (Wald *χ*^*2*^_*1,59*_=11.885, *P*=6×10^−4^), and (*RP/GP*)_i_/(*RP/GP*)_*m*_ and (*PRP/GP*)_i_/(*PRP/GP*)_m_ values smaller (Wald *χ*^*2*^_*1,59*_≥6.77, *P*≤0.009), than one. Finally, in JPN1-TuMV-infected plants *GP*_*i*_*/GP*_*m*_, *RP*_*i*_*/RP*_*m*_ and *PRP*_*i*_*/PRP*_*m*_ did not depend on the allometric group (Wald *χ*^*2*^_*1,166*_≤0.88, *P*≥0.349) (Figure 3F and Supplementary Table S1). Also, (*RP/GP*)_i_/(*RP/GP*)_*m*_ and (*PRP/GP*)_i_/(*PRP/GP*)_m_ did not differ between groups 1 and 2 and showed values smaller than one (Wald *χ* ^*2*^≤0.52, *P*≥0.470) (Figure 3I and Supplementary Table S1). The effect of infection on all plant developmental traits was similar in subgroups 1a and 1b (Wald *χ*^*2*^_*1,166*_≤3.77, *P*≥0.070).

In summary, Arabidopsis fecundity and mortality tolerances to UK1-TuMV are associated with modifications of the plant developmental schedule, whereas fecundity tolerance to LS-CMV and JPN1-TuMV is accompanied by resource reallocation from growth to reproduction. Interestingly, broad-sense heritability of the effect of UK1-TuMV infection on *GP, RP* and *PRP* was higher than that of the effect of infection on *RW* and *IW* (*h*^*2*^ _*b*_ =0.58-0.83 *vs*. 0.40-0.58), whereas the opposite was observed for LS-CMV and JPN1-TuMV infected plants (*h*^*2*^ _*b*_ =0.17-0.39 *vs*. 0.38-0.41 and 0.50-0.61 *vs*. 0.68-0.87, respectively) (Supplementary Table S2). Thus, the plant life-history traits associated with the tolerance response to a given virus have higher host dependency that those not related to tolerance to that particular virus.

### Trade-offs between Arabidopsis defences to virus infection

To analyse Arabidopsis resistance-tolerance trade-offs to each virus, bivariate relationships between virus accumulation and the slope of the *SW* and *LP* to virus accumulation regression were explored, a significantly negative association indicating a trade-off. No significantly negative association was observed between resistance and the two measures of tolerance for any of the three viruses, neither using the whole set of plant genotypes (*r*≤0.23; *P*≥0.367), nor for each allometric group (*r*≤0.40; *P*≥0.223).

We used the same approach to analyse fecundity tolerance-tolerance trade-offs (Figure 4A-C). Bivariate analyses indicated a negative relationship between fecundity tolerance to UK1-TuMV and to LS-CMV (*r*=-0.62; *P*=0.007). No association was found between fecundity tolerance to UK1-TuMV and to JPN1-TuMV (*r*=-0.20; *P*=0.418), but this was just due to the three Subgroup 1a genotypes (*r*=-0.90; *P*<1×10^−4^). Finally, no association was detected between fecundity tolerance to JPN1-TuMV and LS-CMV (*r*=0.25; *P*=0.325), but again removal of Subgroup 1a genotypes resulted in a positive association (*r*=0.65; *P*=0.001) (Figure 4A-C). Because our previous results strongly suggested that trade-offs between tolerance to different viruses were linked to plant allometry, we also analysed such trade-offs by GzLMs virus pairwise comparisons of the slope of the *SW* to virus accumulation regression considering virus and allometric group as factors. A significant interaction was taken as indicative of a tolerance-tolerance trade-off. When fecundity tolerance upon UK1-TuMV infection was compared with that upon infection by the other two viruses, a significant virus per allometric group interaction was observed (Wald *χ* ^*2*^*≥*35.12, *P*<1×10^−4^). On the other hand, the virus genotype per allometric group interaction was not significant when comparing JPN1-TuMV and LS-CMV (Wald *χ*^*2*^_*1,34*_*=*1.87, *P*=0.275) (Figure 2A). Given the bimodal distribution of fecundity tolerance in Group 1 JPN1-TuMV infected plants, we also performed pairwise comparisons considering subgroups 1a and 1b. When fecundity tolerance upon UK1-TuMV and LS-CMV infection was compared between subgroups 1a and 1b, and Group 2, a significant interaction between factors was observed (Wald *χ*^*2*^_*1,28*_*≥*24.89, *P*<1×10^−4^) (Figure 2A). The comparison of JPN1-TuMV and LS-CMV infected plants yielded a significant interaction only between Subgroup 1a and Group 2 (Wald *χ*^*2*^_*1,28*_*=*12.34, *P*=4×10^−4^) (Figures 2A). Conversely, the comparison of UK1-TuMV and JPN1-TuMV infected plants indicated a significant interaction between virus genotype and plant allometry for the combination of Subgroup 1b and Group 2 (Wald *χ*^*2*^_*1,28*_*=*35.97, *P*<1×10^−4^) (Figure 2A). Altogether, these results indicate trade-offs between fecundity tolerance to UK1-TuMV and to the other two viruses.

**Figure 4.**
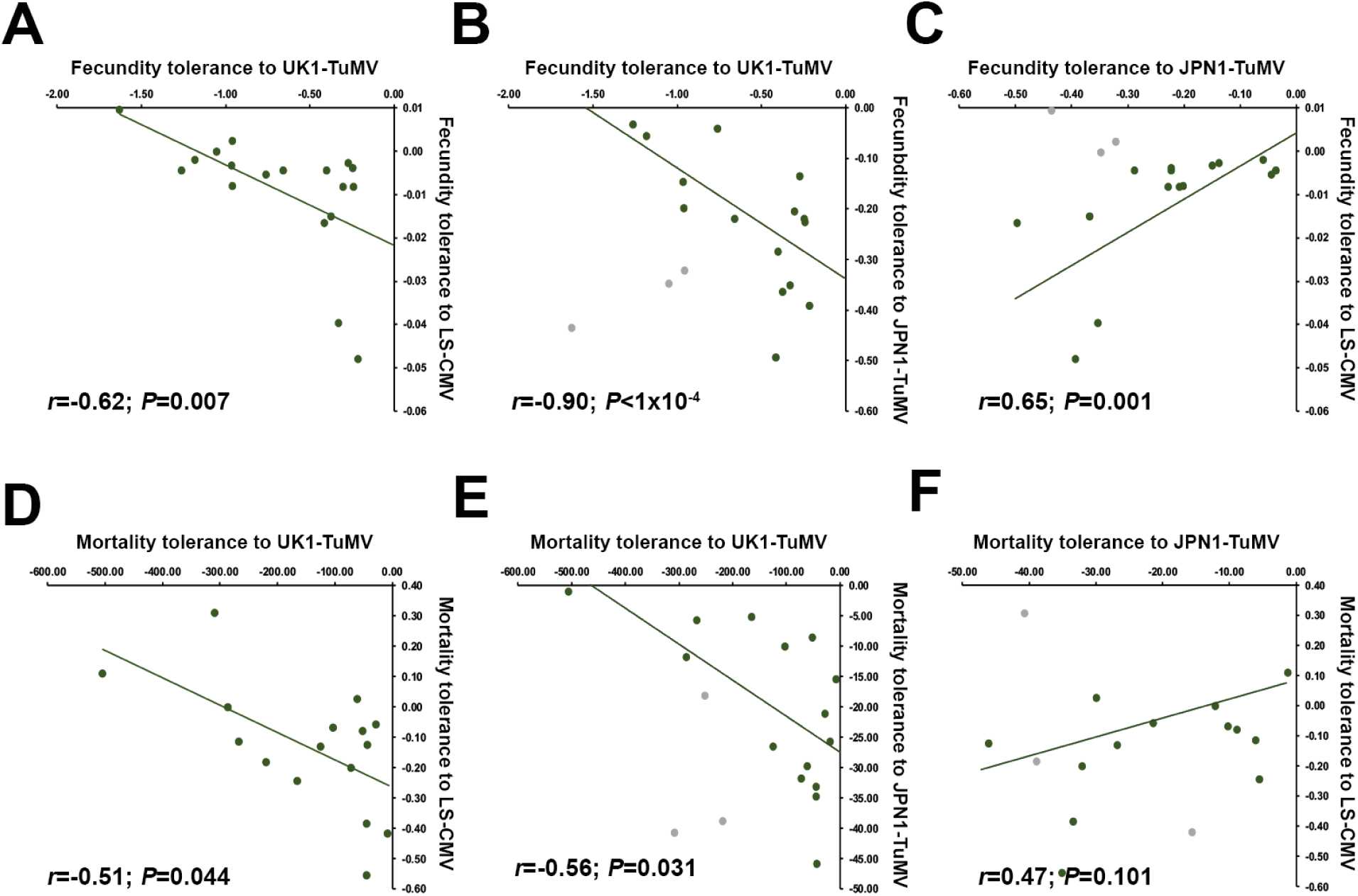
Trade-offs between Arabidopsis tolerances to UK1-TuMV, LS-CMV and JPN1-TuMV. Panels A-C: Pairwise linear regressions between fecundity tolerance to UK1-TuMV, LS-CMV and JPN1-TuMV. Panels D-F: Pairwise linear regressions between mortality tolerance to UK1-TuMV, LS-CMV and JPN1-TuMV. Data are slope of the *SW* (fecundity tolerance) and *LP* (mortality tolerance) to virus accumulation regression for each Arabidopsis genotype. Grey dots correspond to values for Subgroup 1a genotypes, which were excluded from the analyses.

Bivariate analyses indicated a significant negative association between mortality tolerance to UK1-TuMV and to LS-CMV (*r*=-0.51; *P*=0.044), and between tolerance to UK1-TuMV and to JPN1-TuMV when excluding Subgroup 1a genotypes (*r*=-0.56; *P*=0.031). No significant association was observed between mortality tolerance to JPN1-TuMV and to LS-CMV (*r*=0.12; *P*=0.627) even excluding Subgroup 1a genotypes (Figure 4D-F). In addition, the comparison of slope of the *LP* to virus accumulation regression between plants infected by UK1-TuMV and by the other two viruses showed a significant virus per allometric group interaction (Wald *χ* ^*2*^ *≥*29.69, *P*<1×10^−4^), whereas no such interaction was detected between JPN1-TuMV and LS-CMV (Wald *χ*^*2*^_*1,34*_*=*1.26, *P*=0.261) (Figure 2B). When Group 1 plants were divided into Subgroups 1a and 1b, the only significant interaction was between Subgroup 1b and Group 2 genotypes for comparisons of mortality tolerance to UK1-TuMV and JPN1-TuMV (Wald *χ*^*2*^_*1,29*_*=*22.35, *P*<1×10^−4^) (Figure 2B). These results indicate trade-offs between mortality tolerance to UK1-TuMV and to the other two viruses.

For each virus, we also analysed potential mortality-fecundity tolerance trade-offs. No significant bivariate associations were found when considering all plant genotypes together, or each allometric group separately in any of the three viruses (*r*≤0.64; *P*≥0.119). Exception were UK1-TuMV infected plants when analysed as a whole, in which both tolerances showed a positive association (*r*=0.57; *P*=0.013). Thus, when associated, higher mortality tolerance increases fecundity tolerance to a given virus.

## Discussion

Accumulating evidence indicates that tolerance is as widespread as resistance as a plant defence strategy, and therefore central to understand plant-pathogen (including plant-virus) interactions. However, the mechanisms by which tolerance is achieved and the forces shaping its evolution are still poorly understood (*Baucom & de Roode, 2011; Pagán & García-Arenal, 2018*). Using plant-virus interactions, we tested the hypotheses that tolerances to pathogens with different virulence levels are associated with modifications of different plant life-history traits, and that the evolution of tolerance to a given pathogen depends on trade-offs established with the level of tolerance to others.

We showed that Arabidopsis displays genotype-specific fecundity tolerance to the highly virulent virus UK1-TuMV, with plants of the allometric Group 2 having higher tolerance than Group 1 ones. In Arabidopsis, UK1-TuMV infection often prevents seed production (*Sánchez et al., 2015; Vijayan et al., 2017*; this work), such that this virus can be considered as a sterilizing pathogen. Because sterilizing pathogens have an enormous impact on the host fitness, hosts are expected to evolve defences against this type of pathogens (*Lafferty & Kuris, 2009*). Theoretical models on the evolution of host defences predict that infection by a sterilizing pathogen promotes tolerance rather than resistance. Resistance restricts pathogen multiplication and, through repeatedly paying the price to control the pathogen’s growth, resistance would come at infinite cost. Conversely, tolerance would compensate the effect of infection without attempting to control pathogen’s multiplication, thus being less costly (*Restif & Koella, 2004; Best et al., 2010*). Although we did not analyse the costs of resistance and tolerance, our results would support this prediction in that Arabidopsis evolves tolerance to a sterilizing virus rather than resistance: Firstly, half of the Arabidopsis genotypes were not sterilized by UK1-TuMV regardless of virus multiplication, which by definition increases tolerance. Secondly, the level of resistance did not relate with plant fitness, and extreme resistance (immunity) was not detected, indicating that resistance is not associated with the effect of UK1-TuMV on progeny production.

It should be noted that upon UK1-TuMV infection, infected plants of tolerant Arabidopsis genotypes produced on average 30% of the seeds produced by mock individuals. It could be argued that this level of fecundity tolerance is not effective, i.e., seed production of infected plants is far from that of uninfected ones (*Shuckla et al., 2018*). However, mathematical models on the evolution of tolerance to sterilizing pathogens predict that optimal levels of tolerance will not surpass 50% of the progeny produced by uninfected individuals, regardless of tolerance being modelled as a function of host mortality, lifespan or transmission rate (*Restif & Koella, 2004; Hall et al., 2007; Best et al., 2010*). Even if we consider 30% of progeny production upon UK1-TuMV infection as a low level of fecundity tolerance, it would be selectively advantageous for Arabidopsis, as it makes the difference between leaving progeny or not. Indeed, various models showed that this level of fecundity tolerance drives the host population out of the pathogen-driven extinction margins, especially at high levels of pathogen prevalence (*Boots & Sasaki, 2002; Antonovics, 2009*). Accordingly, experimental analyses in other host-sterilizing pathogen interactions reported similar fecundity tolerance levels (*Fredensborg & Poulin, 2006; Vale & Little, 2012*). It is relevant to mention that Arabidopsis fecundity and mortality tolerances to UK1-TuMV were positively associated, whereas upon infection by milder viruses they were not. This observation would agree with models predicting that, for highly virulent parasites, fecundity tolerance is a saturating function of mortality tolerance (*Best et al., 2010*), provided that our data is in the linear part of the curve. Altogether, to our knowledge these results would represent the first example of plant tolerance to a sterilizing virus.

Fecundity tolerance to UK1-TuMV was associated with genotype-specific modifications of the plant developmental schedule. Particularly, upon UK1-TuMV infection more fecundity-tolerant Group 2 genotypes showed shorter growth, and longer reproductive, periods than mock-inoculated plants. This observation agrees with the prediction of the Life-History Theory that bringing forward the age at maturity allows infected hosts to reproduce before they experience the full cost of infection, thus compensating (at least partly) the effect on host fitness (*Hochberg et al., 1992; Gandon et al., 2002*). These results are also in agreement with experimental analyses of life-history modifications upon infection by highly virulent parasites in animals (e.g., *Agnew et al., 2000; Ebert et al., 2004; Fredensborg and Poulin, 2006*). Bringing forward the age at maturity may have important consequences for Arabidopsis population dynamics. Early progeny production would allow seeds from infected plants to germinate and occupy the most suitable niches before uninfected individuals produce theirs, which represents a competitive advantage (*Akiyama & Agren, 2014; Gioria et al., 2018*). This could contribute to compensate the smaller progeny production of infected plants, provided that virus infection does not affect seed viability as shown here. Shorter growth, and longer reproductive, periods of Group 2 genotypes were also associated with higher mortality tolerance to UK1-TuMV. It has been proposed that larger host growth periods caused by pathogen-mediated sterilization allows the storage of reproduction-liberated resources into host growth until the pathogen can exploit them (*Jaenike, 1996; O’Keefe & Antonovics, 2002*). This hypothesis is based on the assumption that host resources can be allocated to either host or pathogen reproduction. Thus, resources dedicated to host reproduction become unavailable for pathogen growth, reducing the effects of infection. This would be the case for the Arabidopsis-UK1-TuMV interaction: Early age at maturity of Group 2 genotypes and subsequent reproduction would reduce the resources available for virus multiplication, limiting/delaying the full cost of infection on plant survival.

Arabidopsis fecundity tolerance to LS-CMV was higher in Group 1 than in Group 2 genotypes, which was associated with resource reallocation from growth to reproduction, an extensively studied response (*Pagán et al. 2007,2008,2009, Hily et al., 2014,2016; Shuckla et al., 2018*). Notably, our results are in agreement with these previous works even if we quantified tolerance as the slope of the fitness to virus load regression rather than at a single pathogen load, and support the Life-History Theory prediction that hosts would evolve tolerance to milder pathogens (as CMV) through resource reallocation from growth to reproduction (*Hochberg et al. 1992; Gandon et al. 2002*). Thus, it could be concluded that Arabidopsis tolerance to plant virus infection is virulence-dependent, which is another prediction of the Life-History Theory. However, our results could be also explained if Arabidopsis life-history trait modifications were virus species-specific, rather than depend on virulence. Indeed, using six Arabidopsis genotypes *Shuckla et al., (2018)* concluded that fecundity tolerance through resource reallocation was specific to CMV, but these authors only considered a highly virulent TuMV isolate. The effect of a milder TuMV genotype (JPN1-TuMV) on Arabidopsis might shed light on this question. Upon JPN1-TuMV infection, half of the Group 1 genotypes showed higher mortality and fecundity tolerances than Group 2 genotypes, all infected plants being fertile, and tolerance being associated with resource reallocation from growth to reproduction. In the other half of Group 1 genotypes, JPN1-TuMV sterilized over 50% of the plants and no tolerance response was observed. Therefore, Arabidopsis Group 1 genotypes in which JPN1-TuMV infection has lower virulence display similar responses to those observed upon LS-CMV infection, whereas in plant genotypes for which JPN1-TuMV virulence is higher the effect of infection resembles to that of UK1-TuMV. This strongly suggests that tolerance is virulence-dependent rather than virus-specific. Note that the subdivision of Group 1 genotypes resulted in 3 to 4 genotypes per subgroup, and the generality of our observations should be validated in a larger number of Arabidopsis genotypes, and in other pathogens and hosts.

We failed in finding a negative association between plant resistance and tolerance to the same virus across Arabidopsis genotypes, which indicates the absence of trade-offs between these two defence mechanisms. On the other hand, Arabidopsis could not optimize at the same time tolerances to viruses displaying different virulence levels (negative association between these tolerances), with LS-CMV and JPN1-TuMV (lower virulence) inducing different and mutually exclusive life-history modifications than UK1-TuMV (higher virulence). A number of experimental works reported that pathogen-driven changes in host life-history traits can be either genetically determined or the consequence of phenotypic plasticity (*Michalakis & Hochberg, 1994; Schlichting & Pigliucci, 1998; McLeod & Day, 2015*). Thus, it could be hypothesized that one or both of these two types of determinisms may be involved in the observed tolerance-tolerance trade-offs. Our data indicates that trade-offs are influenced by two main factors: (i) Virus virulence: Plant genotypes showed different responses in different environments (i.e., virulence levels), which is indicative of phenotypic plasticity (*Michalakis & Hochberg, 1994*). (ii) Plant allometry: Group 1 genotypes showed tolerance to less virulent viruses through resource reallocation, whereas Group 2 genotypes showed tolerance to the most virulent one by altering plant development. Arabidopsis Group 1 genotypes have bigger rosettes and smaller inflorescences than Group 2 ones. That is, in Group 1 genotypes most resources are diverted into growth, whereas in Group 2 resources are primarily dedicated to reproduction. Hence, Group 1 plants would have a relatively wide margin to reallocate growth resources into reproduction; this margin being much narrower, and therefore less efficient, for Group 2 genotypes. In addition, bringing forward the age at maturity requires accelerated rosette growth rates, as Arabidopsis needs to reach a minimum rosette size to flower (*Méndez-Vigo et al., 2010*). Group 1 genotypes typically show faster rosette growth rates (*Hily et al., 2016*), and therefore have less margin to accelerate it, than Group 2 genotypes. Thus, the two allometric groups have particular characteristics that are genetically determined (*Manzano-Piedras et al., 2014*), and that could influence the evolution of tolerance. In support of this genetic determinism, our results indicated that heritability in tolerance-related plant traits was always medium-high. Therefore, although fecundity tolerance is a phenotypically plastic response, the type of response depends on the genetic background of the plant, and tolerance-tolerance trade-offs likely have both genetic and phenotypic plasticity components. This combination of phenotypic plasticity and genetic determinism for tolerance has been also shown in response to other factors such as the moment of plant inoculation, light, temperature and plant density (*Pagán et al. 2007,2009; Hily et al., 2016; Montes & Pagán, 2019*), factors that would modulate the tolerance-tolerance trade-offs observed here, which would be an interesting avenue for future research.

Tolerance-tolerance trade-offs may have important implications for understanding the evolution of host defences. To date, most mathematical models on this subject are built on the assumption that tolerance evolves in single-host-pathogen interactions (*Kutzer & Armitage, 2016; Pagán & García-Arenal, 2018*). These models predict that tolerance would be selectively advantageous for both the host and the pathogen, as tolerance will increase its prevalence, such that genes conferring tolerance will become fixed in the host population (*Rausher, 2001; Råberg et al., 2009*). This is generally applicable to mortality tolerance because it increases the infectious period but would only apply to fecundity tolerance if the pathogen is vertically transmitted (*Best et al., 2008*). In Arabidopsis, CMV and TuMV are seed-transmitted (*Pagán et al. 2014; Montes & Pagán, 2019*). However, our results suggest polymorphisms for both fecundity and mortality tolerance. Increasing evidence indicate that in nature host populations are invaded by more than one pathogen, occurring in single and mixed infections (*Syller, 2012*). Thus, host defences often evolve in a multi-pathogen context. Our results indicate that, in this scenario, the evolution of both fecundity and mortality tolerance to a given virus comes at the cost of higher susceptibility to other(s), which may impose a selection pressure on tolerance and prevent fixation. Hence, more realistic analyses on the evolution of host defences should consider the combined effects of more than one pathogen, and not necessarily in coinfection.

## Materials and methods

### Viruses and Arabidopsis genotypes

Viruses UK1-TuMV (Acc.N. AB194802), JPN1-TuMV (Acc.N. KM094174), and LS-CMV (Acc.N. AF127976) were used. JPN1-TuMV was obtained from a field-infected plant of *Raphanus sativus* (Brassicaceae) and propagated in *Nicotiana benthamiana* plants. UK1-TuMV and LS-CMV were derived from biologically active clones (*Zhang et al., 1994; Sánchez et al., 1998*) by *in vitro* transcription with T7 RNA polymerase (New England Biolabs, Ipswich, USA), and transcripts were used to infect *N. benthamiana* plants for virus multiplication. We used a single CMV isolate because previous analyses indicated that, in Arabidopsis, the fraction of the variance in virulence/tolerance explained by the CMV isolate is very low (4%) (*Pagán et al., 2007*), which is not the case for TuMV. Indeed, UK1-TuMV and JPN1-TuMV have different levels of virulence in Arabidopsis (*Sánchez et al., 2015; Montes & Pagán, 2019*). This allowed exploring whether variation in tolerance to TuMV and CMV were species-specific or virulence-dependent.

We used ten genotypes representing the Eurasian geographic distribution of the species and eight representing its distribution in the Iberian Peninsula, a Pleistocene glacial refuge for Arabidopsis (*Sharbel et al., 2000*) (Table 1). Seeds were stratified for seven days at 4°C in 15cm-diameter pots, 0.43l volume containing 3:1, peat:vermiculite mix. Afterwards, pots were moved for seed germination and plant growth to a greenhouse at 22°C, 16h light (intensity: 120-150 mol s/m^2^), with 65-70% relative humidity. In these conditions, plant genotypes conformed two allometric groups (Table 1 and Supplementary Figure S1) as previously described (*Pagán et al., 2008*). Because plant allometry has been repeatedly reported as a relevant factor to understand Arabidopsis tolerance to virus infection (*Pagán & García-Arenal, 2018*), allometric group was considered as a factor in all analyses. Plants were mechanically inoculated, either with *N. benthamiana* TuMV- and CMV-infected tissue ground in 0.1M Na_2_HPO_4_+0.5M NaH_2_PO_4_+0.02% DIECA, or with inoculation buffer for mock-inoculated plants. Inoculations were done when plants were at developmental stages 1.05-1.06 (*Boyes et al., 2001*). After inoculation, all individuals were randomized in the greenhouse. For each Arabidopsis genotype, seven to ten plants per virus were inoculated, and other seven were mock inoculated.

**Table 1.** Arabidopsis genotypes used in this work, their geographical origin and allometric group/subgroup.

| Genotype | Origin | Allometric group (subgroup) |
| --- | --- | --- |
| Cum-0 | Cumbres Mayores (Spain) | Group 1(a) |
| Kas-0 | Kashmir (India) | Group 1(a) |
| LI-0 | Llagostera (Spain) | Group 1(a) |
| Cad-0 | Candelario (Spain) | Group 1(b) |
| Cdm-0 | Caldas de Miravete (Spain) | Group 1(b) |
| Kas-2 | Kashmir (India) | Group 1(b) |
| Kyo-1 | Kyoto (Japan) | Group 1(b) |
| An-1 | Amberes (Belgium) | Group 2 |
| Bay-0 | Bayreuth (Germany) | Group 2 |
| Col-0 | Columbia (Unknown) | Group 2 |
| Cvi | Cape Verde Islands | Group 2 |
| Fei-0 | Santa María da Feira (Portugal) | Group 2 |
| Ler | Landsberg (Poland) | Group 2 |
| Cen-1 | Centenera (Spain) | Group 2 |
| Mer-0 | Mérida (Spain) | Group 2 |
| Pro-0 | Proaza (Spain) | Group 2 |
| Shak | Shakdara (Tadjikistan) | Group 2 |
| Ver-5 | Verin (Spain) | Group 2 |

### Quantification of virus multiplication

Virus multiplication was quantified as viral RNA accumulation 15 days post-inoculation via qRT-PCR and was used as a measure of plant resistance to virus infection. For each plant, four leaf disks of 4mm in diameter from four systemically-infected rosette leaves were collected. Total RNA extracts were obtained using TRIzol^®^ reagent (Life Technologies, Carlsbad, USA), and 0.32ng of total RNA were added to the Brilliant III Ultra-Fast SYBR Green qRT-PCR Master Mix (Agilent Technologies, Santa Clara, USA) according to manufacturer’s recommendations. Specific primers were used to amplify a 70nt fragment of the TuMV, and a 106nt fragment of the CMV, coat protein (CP) gene, respectively (*Lunello et al., 2007; Hily et al., 2014*). Each sample was assayed by triplicate on a Light Cycler 480 II real-time PCR system (Roche, Indianapolis, USA). Absolute viral RNA accumulation was quantified as ng of viral RNA/μg of total RNA utilizing internal standards. For the two TuMV isolates, internal standards consisted in ten-fold dilution series of plasmid-derived RNA transcripts of the same 70nt CP fragment from UK1-TuMV. For LS-CMV, ten-fold dilution series were prepared using purified viral RNA. Internal standards ranged from 2×10^−3^ng to 2×10^−7^ng.

### Effect of infection on plant growth and reproduction

Aboveground plant structures were harvested at complete senescence. The weights of the rosette (*RW*), inflorescence (*IW*), and seeds (*SW*) were obtained. *RW* was used to estimate plant resources dedicated to growth, and *IW* and *SW* were utilized to estimate plant resources dedicated to reproduction (*Thompson & Stewart, 1981*). The effect of virus infection on these traits was quantified by calculating infected to mock-inoculated plants ratios for each of them, dividing the value of each infected plant by the mean value of the mock-inoculated plants of the same genotype (*Trait*_*i*_*/Trait*_*m*_, *i* and *m* denoting infected and mock-inoculated plants, respectively). Following *Pagán et al., (2008)*, resource reallocation from growth to reproduction upon virus infection was analysed by calculating *(IW/RW)*_*i*_*/(IW/RW)*_*m*_ and *(SW/RW)*_*i*_*/(SW/RW)*_*m*_ ratios. Values of these ratios greater than one were considered as indicative of such resource reallocation. Seed viability, estimated as per cent germination, did not significantly differ between mock-inoculated (93.0-99.3%) and infected (91.0-99.7%) plants (*χ*^*2*^≤2.16; *P*≥0.096). Also, virus infection did not affect the weight of a single seed (Wald *χ*^*2*^≤0.99; *P*≥0.110) (Supplementary Table 3). Thus, *SW* similarly reflects the number of viable seeds in both mock-inoculated and infected plants.

### Effect of infection on plant development

We recorded growth period (*GP*), as days from inoculation to the opening of the first flower; reproductive period (*RP*), as days from the opening of the first flower to the shattering of the first silique; and plant post-reproductive period (*PRP*), as days from the shattering of the first silique to plant senescence. In Arabidopsis, the opening of the first flower co-occurs with the end of the rosette growth, and the shattering of the first silique co-occurs with the end of flower production (*Boyes et al., 2001*). The total life period (*LP*) was quantified as the sum of the three periods. The effect of virus infection on *GP, RP* and *PRP*, was quantified as infected to mock-inoculated plants ratios. The *(RP/GP)*_*i*_*/(RP/GP)*_*m*_ and (*PRP/GP*)_*i*_*/(PRP/GP)*_*m*_ ratios were used to analyse virus-induced alterations of plant development.

### Tolerance measure

Following *Little et al., (2010)* and *Råberg (2014)*, range fecundity and mortality tolerances of each Arabidopsis genotype were calculated as the slope of the linear regression of *SW* and *LP*, respectively, to virus accumulation considering both infected and mock-inoculated plants.

### Statistical analysis

Analysed traits were not normally distributed, and variances were heterogeneous. Therefore, differences between viruses, plant genotypes and allometric groups/subgroups were analysed by Generalized Linear Mixed Models (GzLMMs) considering virus as fixed factor, and Arabidopsis genotype as random factor, which was nested to allometric group/subgroup (considered as fixed factor). Trade-offs between resistance, fecundity tolerance and mortality tolerance were analysed using Spearman’s test. Tolerance-tolerance trade-offs according to virus and plant allometric group/subgroup were analysed using Generalized Linear Models (GzLMs), considering both as fixed factors. Broad-sense heritability was estimated as *h* ^*2*^ _*b*_ *=V* _*G*_ */(V* _*G*_ *+V* _*E*_ ), where *V* _*G*_ is the among-genotypes variance component and *V*_*E*_ is the residual variance. Variance components were determined using GzLMMs by the REML method (*Lynch & Walsh, 1998*). GzLMMs and GzLMs were performed using R-libraries lme4, nlme and lmerTest (*Douglas et al., 2015, Kuznetsova et al., 2017, Pinheiro et al., 2018*). Statistical analyses were conducted using R version 3.5.0 (*R Core Team, 2018*).

## Acknowledgements

Marisa López-Herranz and Miriam Gil-Valle provided excellent technical assistance. This work was supported by Plan Nacional I+D+i, MINECO, Spain [BIO2016-79165-R] to IP. NM was supported by Ministerio de Economía y Competitividad (Instituto de Salud Carlos III) [PIE13/00041], and VV by an Erasmus Mundus BRAVE Scholarship of the EU [2013-2536/001-001].

